# Multiple paths towards repeated phenotypic evolution in the spiny-leg adaptive radiation (*Tetragnatha*; Hawaii)

**DOI:** 10.1101/2022.11.29.518358

**Authors:** José Cerca, Darko D. Cotoras, Cindy G. Santander, Vanessa C. Bieker, Leke Hutchins, Jaime Morin-Lagos, Carlos F. Prada, Susan Kennedy, Henrik Krehenwinkel, Andrew J. Rominger, Joana Meier, Dimitar Dimitrov, Torsten H. Struck, Rosemary G. Gillespie

## Abstract

The repeated evolution of phenotypes is ubiquitous in nature and offers some of the clearest evidence of the role of natural selection in evolution. The genomic basis of repeated phenotypic evolution is often complex and can arise from a combination of gene flow, shared ancestral polymorphism and *de novo* mutation. Here, we investigate the genomic basis of repeated ecomorph evolution in the adaptive radiation of the Hawaiian spiny-leg *Tetragnatha*. This radiation comprises four ecomorphs that are microhabitat-specialists, and differ in body pigmentation and size (Green, Large Brown, Maroon, and Small Brown). Using 76 newly generated low-coverage, whole-genome resequencing samples, coupled with population genomic and phylogenomic tools, we studied the evolutionary history of the radiation to understand the evolution of the spiny-leg lineage and the genetic underpinnings of ecomorph evolution. Congruent with previous works, we find that each ecomorph has evolved twice, with the exception of the Small Brown ecomorph, which has evolved three times. The evolution of the Maroon and the Small Brown ecomorphs likely involved ancestral hybridization events, whereas the Green and the Large Brown ecomorphs likely evolved because of either standing genetic variation or *de novo* mutation. Pairwise comparisons of ecomorphs based on the fixation index (F_ST_) show that divergent genomic regions include genes with functions associated with pigmentation (melanization), learning, neuronal and synapse activity, and circadian rhythms. These results show that the repeated evolution of ecomorphs in the Hawaiian spiny-leg *Tetragnatha* is linked to multiple genomic regions and suggests a previously unknown role of learning and circadian rhythms in ecomorph.

## Introduction

Adaptive radiation, the evolutionary process where an ancestral lineage diversifies into multiple phenotypically-distinct species which occupy different ecological niches, offers a natural experiment to disentangle links between the phenotypic diversification and environmental adaptation (Schluter 2000; Gillespie et al. 2020). Of particular interest in adaptive radiation is the repeated phenotypic evolution (i.e. parallel or convergent phenotypic evolution) (Cerca 2022), where equivalent phenotypes evolve in response to similar ecological challenges (Losos and Ricklefs 2009; Losos 2010). Repeated evolution of similar phenotypes has been found in multiple radiations and offers a powerful approach for disentangling recurrent and potentially-deterministic phenotypic outcomes in response to similar environmental conditions (Losos 2010, 2011; Gillespie et al. 2018, 2020; Malinsky et al. 2018; Salzburger 2018; Masonick et al. 2022; Urban et al. 2022). For instance, the repeated evolution of habitat-specialists, termed ecomorphs, has been reported in multiple adaptive radiations including the Caribbean *Anolis* lizards (Losos and Ricklefs 2009) and Hawaiian *Tetragnatha* (Gillespie 2004) and *Ariamnes* (Gillespie et al. 2018) spiders. In these cases, the repeated evolution of ecomorphs has been explained based on the spatial segregation of environments, as different islands offer similar environmental conditions (Losos and Ricklefs 2009; Losos 2010). However, despite the deluge of genomic data, and new insights into phenomena of admixture, epigenetics, and other phenomena that highlight the overall flexibility of the genome, we know little about the genomic underpinnings involved in the evolution of discrete phenotypes that appear to arise repeatedly in response to similar selective pressures. The current study set out to examine the genomic basis of recurrent ecomorph evolution in a lineage of Hawaiian spiders.

The Hawaiian *Tetragnatha* spiny-leg species belong to a clade comprising ∼17 species, and are part of a large radiation endemic to the archipelago (Gillespie 2016; Kennedy et al. 2022). These species can be grouped into four ecomorphs (Figure 1 A-D), which are linked to the substrate they inhabit (Gillespie 2004): the Large Brown ecomorph is found on tree bark (Figure 1A), the Green ecomorph on leaves (Figure 1B), the Maroon ecomorph on mosses (Figure 1C), and the Small Brown ecomorph on twigs (Figure 1D). Ancestral character-state reconstructions suggest that the Green ecomorph is likely the ancestral form, having evolved once, while the remaining ecomorphs each evolved twice (Gillespie 2004). Recent genomic work has shown that co-occurring closely related species belonging to the Green ecomorph do not hybridize, and it has been argued that there may be some overlap in their ecological niches in the early stages of diversification, suggesting a possible avenue for the divergence of ecomorphs through character displacement upon secondary contact (Schluter 2000; Cotoras et al. 2018). The genomic bases of ecomorph evolution, which will allow for the ecological differentiation of these closely related and ecologically equivalent species are still poorly understood.

**Figure 1.**
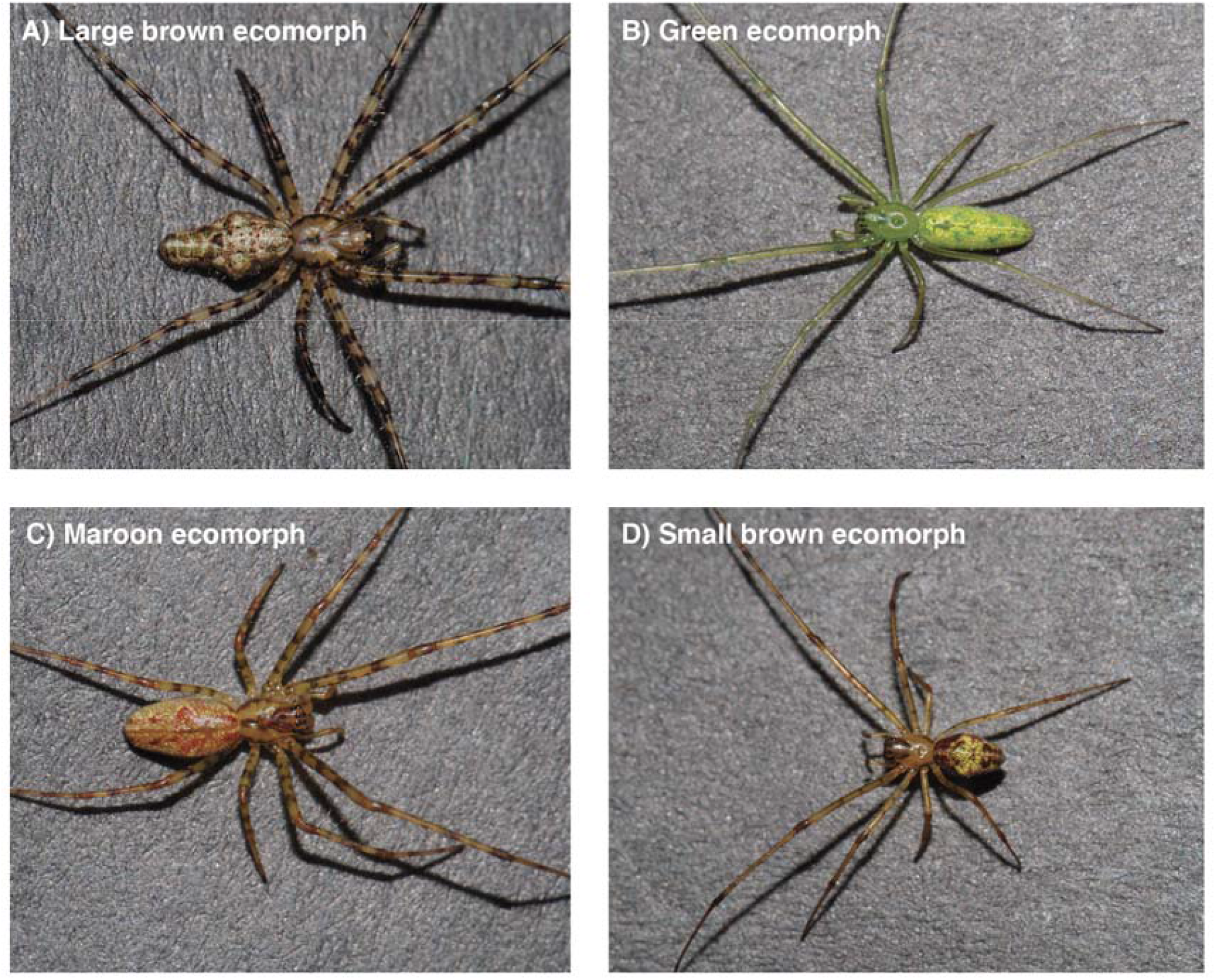
***Tetragnatha* spiny-leg ecomorphs** including representatives of (A) the Large Brown (*Tetragnatha quasimodo*), (B) the Green (*Tetragnatha brevignatha*), (C) the Maroon (*Tetragnatha kamakou*), and (D) the Small Brown (Tetragnatha anuenue). Photographs by Darko D. Cotoras.

The repeated evolution of a phenotype can arise as a result of three non-mutually exclusive genomic processes (Stern 2013; Pease et al. 2016; Lee and Coop 2017, 2019): *de novo* mutation (different mutations causing similar phenotypes), shared ancestral polymorphism (standing variation, old genetic variation being re-recruited), or hybridization (where a particular allele is recruited from one lineage to another). All three patterns have been observed in the context of adaptive radiations (Meier et al. 2018; Choi et al. 2021; Sowersby et al. 2021; De-Kayne et al. 2022). Because they leave distinct footprints along the genome, genomic-level data is able to tease apart the contribution of each of these processes (Lee and Coop 2017).

Here, we whole-genome re-sequence 76 genomes across the *Tetragnatha* spiny-leg radiation with the aim of understanding the genomic basis of repeated ecomorph evolution. We hypothesise that standing genetic variation has been the source for phenotypic repetition in *Tetragnatha* spiny-legs. We start by reconstructing the evolutionary history of the spiny-leg lineage using phylogenetic tools. Then, we explore patterns of excess allele sharing to infer potential hybridization. Finally, we investigate genomic divergence (F_ST_) to understand differentiation between ecomorphs. Our results uncover a complex evolutionary history, showing that repeated evolution may have emerged through multiple different genomic processes.

## Methods

### Field collection

Spiny-leg *Tetragnatha* live in the montane rain forests of all the major Hawaiian islands (1,200-1,800 meters) (Roderick et al. 2012). Specimens were collected either by hand (at night, when spiders are active) or using a beating sheet (both day and night). Specimens were preserved in 95% ethanol and stored at -20°C. A list of specimens including their volcano and island is found in Supplementary Table 01.

### Molecular data generation

We sequenced genomes from a total of 76 individuals (Supplementary Table 01). For each specimen, we extracted DNA from 2-4 legs. Legs were first ground using a tube pestle and incubated overnight in a solution of lysis buffer (10mM Tris pH, 100mM NaCl, 10mM EDTA, 0.5% SDS) and proteinase K at 54° C, we then followed the instructions of the commercial provider Qiagen to extract the DNA. We diluted the DNA in 50 μl of elution buffer and assessed the DNA concentration of each of the 76 extracts using a Qubit fluorometer (ThermoFisher).

Samples with more than 500 ng of DNA were submitted to the QB3-Berkeley, Vincent J. Coates Genomics Sequencing Laboratory at UC Berkeley, where library preparation was conducted. DNA was fragmented using a Bioruptor Pico (Diagenode) and libraries prepared using the KAPA Hyper Prep kit for DNA (KK8504). This involved adding truncated universal stub adapters for DNA-adapter ligation, and indexed primers for PCR amplification (to complete the adapters). The quality of the samples was checked on an AATI (now Agilent) Fragment Analyzer, and the molarity of the library was measured using quantitative PCR with the KAPA Library Quantification Kit (Roche KK4824) on a BioRad CFX Connect thermal cycler. Libraries were then pooled by molarity and sequenced on an Illumina NovaSeq 6000 S4 flowcell for 2 × 150 cycles, with the aim of obtaining 10 Gbp per sample (∼9.5x sequencing depth of the genome). Raw sequencing data was converted to FASTQ format and demultiplexed using the Illumina bcl2fastq2 software while allowing for up to one mismatch in the index sequences.

### Bioinformatics data processing

All of the steps below are reported in GITHUB JOSE. We checked the quality of the sequencing data using fastQC v0.11.8 (Andrews 2017), and identified adapters using AdapterRemoval v2.3.1 (--identify-adapters) (Schubert et al. 2016). We then used Trimmomatic v0.39 (Bolger et al. 2014) to remove adapters and poor quality reads by specifying: maximum 2 mismatches in the adapter sequence; cut three base pairs in the beginning and the end of the read if the quality drops below 20 in these regions; a sliding window of 4 bp with a minimum quality threshold of 20, and a minimum read length of 50. These high-quality reads were aligned to the *Tetragnatha kauaiensis* reference genome v1 (∼1.1 Gb) (Cerca et al. 2021a) using the Burrows-Wheeler Aligner v0.7.17, mem algorithm (Li and Durbin 2009). Because Trimmomatic separates reads as forward-paired, reverse-paired, forward-unpaired, and reverse-unpaired, we aligned all four files to the reference genome, and merged and sorted the final file using Samtools v1.10 (Li et al. 2009). We then estimated mapping quality of each alignment using samtools flagstat, finding no mapping-bias (Supplementary Table 01), and marked duplicates using GATK 4.1.4.0 (McKenna et al. 2010), by running the SortSam algorithm which sorted reads based on genomic coordinates, followed by the MarkDuplicates function. We then built indexes for each alignment using BuildBamIndex (McKenna et al. 2010), and filtered reads with a low mapping quality (MAPQ) using samtools. Specifically, we discarded reads with mapping quality below 30, and calculated the sequencing depth using samtools depth (option -a was used to output all sequencing positions, even those with 0 depth; Supplementary Table 01). We then processed the data using ANGSD v0.935 (Kousathanas et al. 2017), a pipeline designed to handle and analyze low coverage sequencing data.

### Phylogenetic analyses

We started by doing a phylogenetic reconstruction while accounting for low-coverage to understand the ancestral relationships of ecomorphs. As part of this analysis, we included five specimens from four species belonging to an outgroup lineage, the *Tetragnatha* web builder radiation, which is also endemic to Hawai’i (*Tetragnatha maka, Tetragnatha acuta, Tetragnatha filiciphilia*, and *Tetragnatha stelarobusta*). We started by calling variants in ANGSD using the GATK genotype likelihood model to output a beagle file (-GL 2 -doGlf 2), specifying that a variant should be present in at least 38 individuals (half of the dataset), minimum base quality of 30 (-minQ), a *p*-value threshold of 1e-6 (-SNP_pval), to remove allele counts in less than 5% of the dataset (-minMaf 0.05), and to infer major and minor alleles from the genotype likelihoods (-doMajorMinor 1). The output beagle file was then processed using an in-house script to remove variants in repeat regions by downloading the general feature format file (gff3) from (Cerca et al. 2021a), including the genomic location of repeat elements, converting it to a bed file format using bedtools v2.26.0 (Quinlan and Hall 2010), and generating a white-list of the genome (i.e. non-repeated genomic areas) using an in-house script (See Github). This dataset was used as input in NgsDist (Vieira et al. 2015), where we specified a block size of 20 SNPs, and 100 bootstrap replicates. We then ran the NJ software fastme v2 (Lefort et al. 2015), merged all the 100 replicates into a final tree with bootstrap support using RAXML while specifying an optimization of branch-length, and specifying GTRCAT (GTR + Optimization of substitution rates + Optimization of site-specific) as the substitution model.

Because different parts of the genome can have different topologies as a result of different evolutionary mechanisms, we additionally performed a mitochondrial phylogenetic reconstruction and a phylogenetic reconstruction based on Ultra Conserved Elements (UCE). We retrieved mitochondrial genomes and did a tree reconstruction to analyse nuclear-mitochondrial discordances; which are typically associated with hybridization events. We started by extracting the mitochondrial genomes from the cleaned Illumina libraries using Novoplasty v4.2 (Dierckxsens et al. 2017), a method which uses an iterative baiting-and-expand approach to construct genomic regions. We obtained a collection of seeds encompassing different mitochondrial genes (16S rDNA (hereafter 16S), Cytochrome Oxidase I (COI), and Cytochrome B (CytB)) for species from the radiation (Kennedy et al. 2022). For each specimen, we ran all seeds until we obtained a circularized mitochondrial genome. If no complete mitochondrial genome was obtained for a given specimen, we took the longest contig obtained by Novoplasty. In total, we obtained 46 complete fully-circularized mitochondrial genomes, and 19 partially complete mitochondrial genomes (>5,000 bp). We then concatenated and aligned the 65 mitogenomes using Mafft v7.130b (Katoh and Standley 2013), trimmed the ends of the alignments and ran a tree using IQ-Tree v2.0.3 specifying 1,000 ultrabootstrap replicates and automatic determination of substitution models.

We obtained a UCE dataset to complement the ngsDist methods with a maximum likelihood-based analysis. For each sample, we started by calling every position of the genome using the Bcftools’ v1.10.2 mpileup and call algorithms (Danecek et al. 2021), using the *T. kauaiensis* genome v1 as reference. The output files were normalized for indels using bcftools norm, and filtered for gaps bigger than 4 base pairs, base quality above 20, depth above 1 and below 30. These files were then turned into consensus fasta sequences using bcftools consensus. To extract UCEs from these consensus files, we used the phyluce pipeline (Faircloth 2016), with the ‘Arachnida-UCE-1.1K-v1’ set as bait. Phyluce allows extraction of UCEs, but because of the missing data due to low coverage and fragmented genomes, we were only able to retrieve 29 UCEs that were present in half of the dataset (16 of the UCEs were present in all individuals; and 23 UCEs present in ≥70 individuals).

### Partitioning of genetic variation and hybridization

After reconstructing species and ecomorph relationships we analysed the partitioning of genetic variation and whether hybridization has occurred in the radiation. We studied the partitioning of genetic variation by running a NGSAdmix analysis. This analysis required processing the data with ANGSD, similar to the one used for the phylogenetic reconstruction with NgsDist, but with three exceptions. Specifically, we included a minimum base quality of 20 (-minQ) filter, removed variants with a minimum allele frequency of 0.05 (-minMaf 0.05), and inferred major and minor alleles (-doMajorMinor 1). After filtering for variants in repeat regions, we filtered the data for severe deviations from the Hardy-Weinberg equilibrium (HWE) using PCAngsd, which estimated a likelihood ratio test (LTR) to identify sites where severe deviations of the HWE occur (LTR >24, following recommendations from the contributors of ANGSD; https://github.com/GenisGE/grantsGazelleScripts) (Garcia-Erill et al. 2021). Additionally, given that admixture-like analyses assume that variants are independent, we implemented a linkage disequilibrium (LD) filter. This was done by calculating LD in windows of 100,000 bps using plink (Purcell et al. 2007) and using an in-house script explore LD-decay. We LD-pruned the data by removing genomic regions where linkage, measured as R2, was above 0.11 in windows of 25,000 bps (5,000 bp steps). We then ran NGSadmix analyses (Skotte et al. 2013). We specified cluster values (*K*) between 1-20, and for each we performed 10 independent runs until each reached chain convergence. We estimated the best *K* to be 2 using the Evanno method (Evanno et al. 2005) as implemented as part of CLUMPAK (Kopelman et al. 2015). We present the results for *K*=2, 3, and 15 as ADMIXTURE plots help to clarify how genetic variation is partitioned at different levels (Meirmans 2015).

Considering the nuclear-mitochondrial discordance, we calculated excess of shared alleles in the dataset. To do so, we re-ran ANGSD, with the parameter specification as adone for the NGSadmix analyses, further specifying a genotype likelihood depth filter between 7 and 30 (-geno_minDepth 7 -geno_maxDepth 30). The outputted genotype file was processed to remove variants in repeat regions and the excess allele sharing was calculated using Patterson’s D (ABBA-BABA) and *F*4 ratios (Patterson et al. 2012) using Dsuite (Malinsky et al. 2021). Both these analyses benefited from a tree-backbone and we specified the tree obtained in ngsDist.

### Scans of selection

Whole-genome level data allows understanding whether certain genes or genomic regions are under selection as this process leads to shifts in allelic frequencies (Ravinet et al. 2017). In order to calculate population differentiation (F_ST_), we obtained the Site Frequency Spectrum (SFS) using ANGSD. As our goal is to understand the evolution of ecomorphs, we analysed the data for each monophyletic ecomorph group, based on the nuclear phylogenetic reconstruction obtained with NGSdist. Furthermore, because SFS and F_ST_ estimations require allelic frequencies from multiple individuals, we selected only ecomorphs with 5 or more specimens (Supplementary Table 01). To obtain SFS estimates we ran ANGSD separately for: *Tetragnatha pilosa* (Large Brown group A), *T. quasimodo* (Large Brown group B), *T. kauaiensis* (Green group A), *T. tantalus, T. polychromata, T. brevignatha, T. waikamoi* (Green group B), *T. obscura, T. kukuiki, T. kikokiko, T. anuenue* (Small Brown ecomorph), and *T. kamakou* (Maroon ecomorph). We ran ANGSD independently for each of these groups, specifying that loci had to be in at least 5 individuals, keeping only reads with a single mapping (-uniqueOnly), discarding bad reads (-remove_bads), and keeping variants with a minimum base quality of 20. This yielded a 1-dimensional SFS for each population, which was then processed using realSFS (using -fold 1) to obtain 2-dimensional SFSs for the following pairs: the two Large Browns (Large Brown A vs Large Brown B), the two Greens (Green A vs Green B), Green vs Maroon (Green B vs Maroon), and Small Brown vs Large Brown (Small Brown vs LargeBrown B). For each of the 2-dimensional SFS, we ran the realSFS F_ST_ index algorithm, followed by realSFS F_ST_ stat to obtain the overall F_ST_. Because we were interested in identifying genomic areas of divergence and loci within, we ran the realSFS F_ST_ stat2 algorithm to obtain F_ST_ in 5,000 bp non-overlaping windows. We identified regions of differentiation by Z-transferring F_ST_ and using a Z cut-off of >3. After identifying significantly differentiated regions, we retrieved genes for these windows from the gene annotations of the *T. kauaiensis* genome (Cerca et al. 2021a). We then called orthologs between *T. kauaiensis* and the *Drosophila melanogaster* genome annotations to obtain evidence on gene function, benefiting from the decades of functional genetic research on the latter, using *OrthoFinder2* (Emms and Kelly 2015). After identifying the closest *D. melanogaster* ortholog for a particular gene on a region with high differentiation, we reviewed and summarized the literature on that gene in Supplementary Table 2. When possible, we read at least three papers for each gene.

### Hybridization along the genome

We found evidence for hybridization, and because this process can leave tracks along the genome, we performed an analyses to understand the breadth and impact of hybridization on the genomes. Specifically, the results showed a strong signal of admixture between *T. kamakou* (Maroon ecomorph), *T. perreirai* (Maroon ecomorph), and *T. restricta* (light Brown ecomorph) and to understand whether hybridization occurred we did a TWISST analysis on the three biggest scaffolds and D-suite investigate for the scaffolds with melanization genes. For the TWISST analysis, we selected *T. anuenue, T. perreirai, T. restricta*, and *T. mohihi*. We started by running ANGSD for non-overlapping regions of 5kb segments of the genome, specifying a minimum map quality of 30, minimum base quality of 20, outputting the frequency of the different bases. We then did a NJ-tree for each of these regions using the R package ape (Paradis et al. 2008). We then ran TWISST specifying *T. anuenue*. For the windowed Patterson’s D, we used Dsuite investigate on the scaffolds with melanization genes.

## Results

### Tree reconstruction

In the nuclear tree, *T. pilosa* is sister to the remaining spiny-leg species. The node including all other spiny-legs, however, has a bootstrap support below 95, while all other interspecific nodes have a bootstrap support of 100. Furthermore, the node including all spiny-legs but *T. pilosa* is preceded by a very short branch, and separates two clades: one clade including *T. mohihi* and *T. kauaiensis*, and another clade with all remaining species. The remaining species can be separated in two major clades, the first (clade B) including the Large Brown *T. quasimodo* and a group of Small Brown species including *T. obscura, T. kukuiki, T. kikokiko, T. anuenue*; and the second (clade C) including a group of species representing the Green ecomorph (*T. tantalus, T. polychromata, T. waikamoi, T. brevignatha*), which is sister to a group comprising two Maroon species (*T. perreirai* and *T. kamakou*) and a Small Brown species (*T. restricta*). The maximum-likelihood UCE tree is topologically concordant with the ngsDist tree at the ecomorph level (Supplementary Figure 01). The only topological discordance is the species’ placement within the Small Brown group as part of clade B (*T. obscura, T. kukuiki, T. kikokiko, T. anuenue*). Specifically, the ngsDist tree (Figure 2) shows *T. kukuiki* as sister to *T. obscura*, and *T. kikokiko* as sister to *T. anuenue*, whereas on the UCE tree, *T. anuenue* is sister to a clade including all the aforementioned species (Supplementary Figure 01).

**Figure 2.**
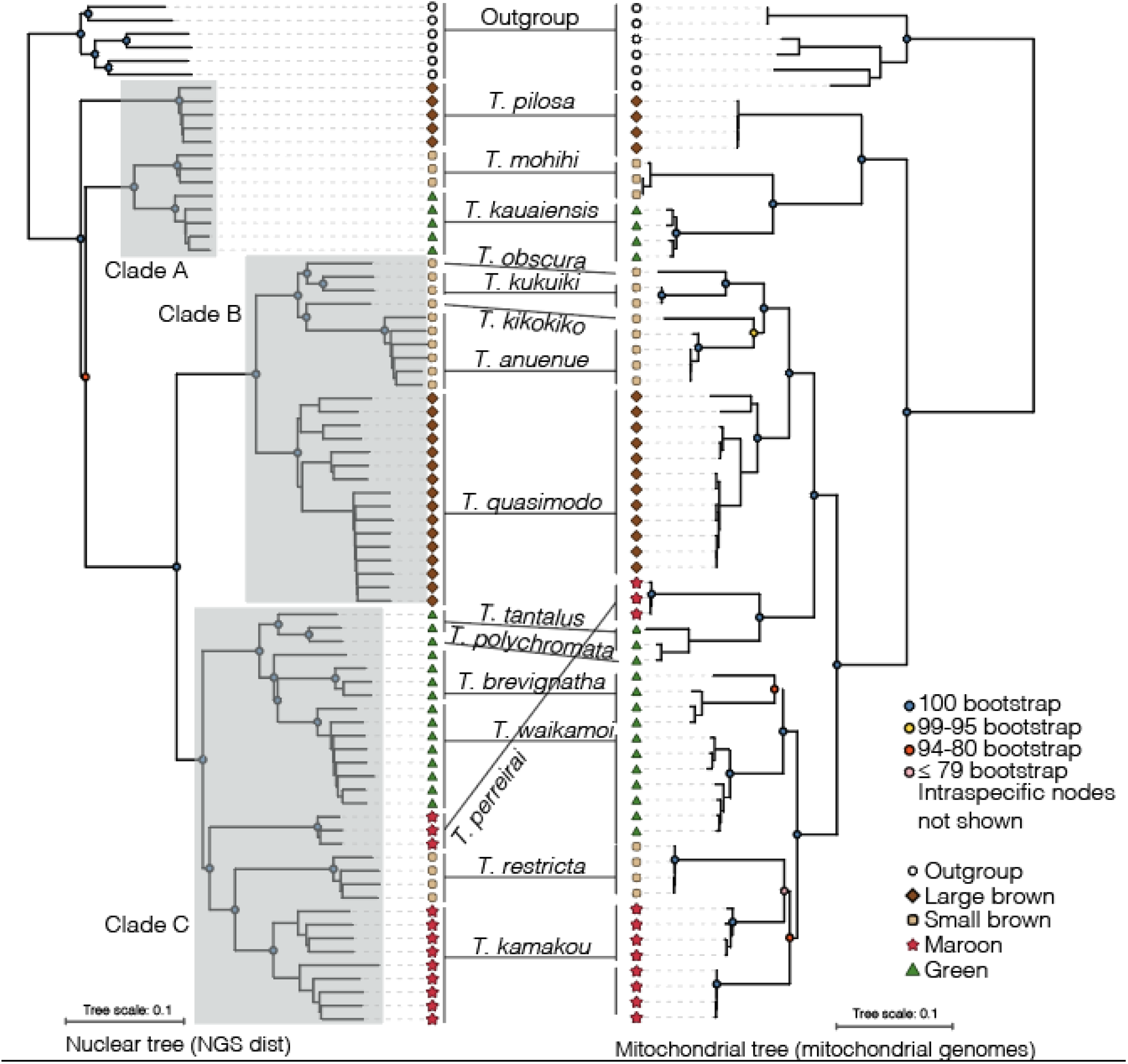
Nuclear and mitochondrial tree reconstructions. The nuclear tree (left) obtained by NgsDist is based on 1,848,915 variable sites. The mitochondrial tree (right) obtained by NovoPlasty and IQ-Tree is based on complete and partial mitochondrial genomes. Bootstrap is provided for each inter-specific node, and the tips of the phylogeny include ecomorph and outgroup labelling as coloured symbols. Clades A-C are plotted in the nuclear tree.

There are mitochondrial-nuclear tree discordances based on topology and bootstrap support (Figure 2). First, in the mitochondrial tree, *T. pilosa* groups together with *T. mohihi* and *T. kauaiensis*, forming a clade while *T. pilosa* branches separately in the nuclear tree. Notably, all basal interspecific nodes have a bootstrap support of 100. Second, *T. tantalus* and *T. polychromata* (Green ecomorphs) are sister to *T. perreirai* (Maroon ecomorph) in the mitochondrial tree, and the clade comprising these three species is sister to the clade including the Large Brown (*T. quasimodo*) and the Small Brown ecomorphs (*T. obscura, T. kukuiki, T. kikokiko, T. anuenue*). These also have 100 bootstrap support at the relevant interspecific nodes. The mitochondrial tree shows that the evolution of Green ecomorphs has occurred three times (1 - *T. kauaiensis;* 2 - *T. tantalus* and *T. polychromata;* 3 - *T. waikamoi* and *T. brevignatha*), as opposed to two clades in the nuclear tree (1 - *T. kauaiensis;* 2 - *T. tantalus, T. polychromata, T. waikamoi* and *T. brevignatha*). Third, the mitochondrial dataset shows *T. kamakou* as paraphyletic, separating the populations from Moloka⍰i and Maui (East Maui volcano). However, this node is weakly supported as it has a bootstrap value below 80.

For simplicity, and based on the mitochondrial phylogenetic reconstruction (and ngsAdmix below), we refer to clades A (*T. kauaiensis, T. pilosa, T. mohihi*), B (*T. anuenue, T. obscura, T. kukuiki, T. kikokiko, T. quasimodo*) and C (*T. kamakou, T. restricta, T. perreirai, T. tantalus, T. polychromata, T. waikamoi and T. brevignatha*) hereafter.

### Structuring of genetic diversity

The best *K* obtained by the Evanno method was *K* = 2 (Figure 3). This *K* separated clade B from the remaining clades (A and C), while *K* = 3 separated all three clades obtained in the phylogeny. There is evidence of a shared history for both *K* values, some species having minor components of other clades. For instance, *T. kikokiko* (Clade B) has a minor component of ancestry from clades A-C on *K* = 2, and a minor component from clade C on *K* = 3. We also display *K* = 15 because that is the number of species in the dataset (Figure 3). For K = 15, some species are assigned to the same genetic cluster: *T. kauaiensis and T. mohihi* (red), *T. obscura and T. kukuiki* (green), and *T. polychromata* and *T. tantalus* (aqua-blue). Two species, interestingly both Large Browns, have intra-specific population structure (*T. pilosa* has two colours, while *T. quasimodo* has three). Only one *T. brevignatha* sample has mixed ancestries, all in common with species belonging to the Green ecomorph.

**Figure 3.**
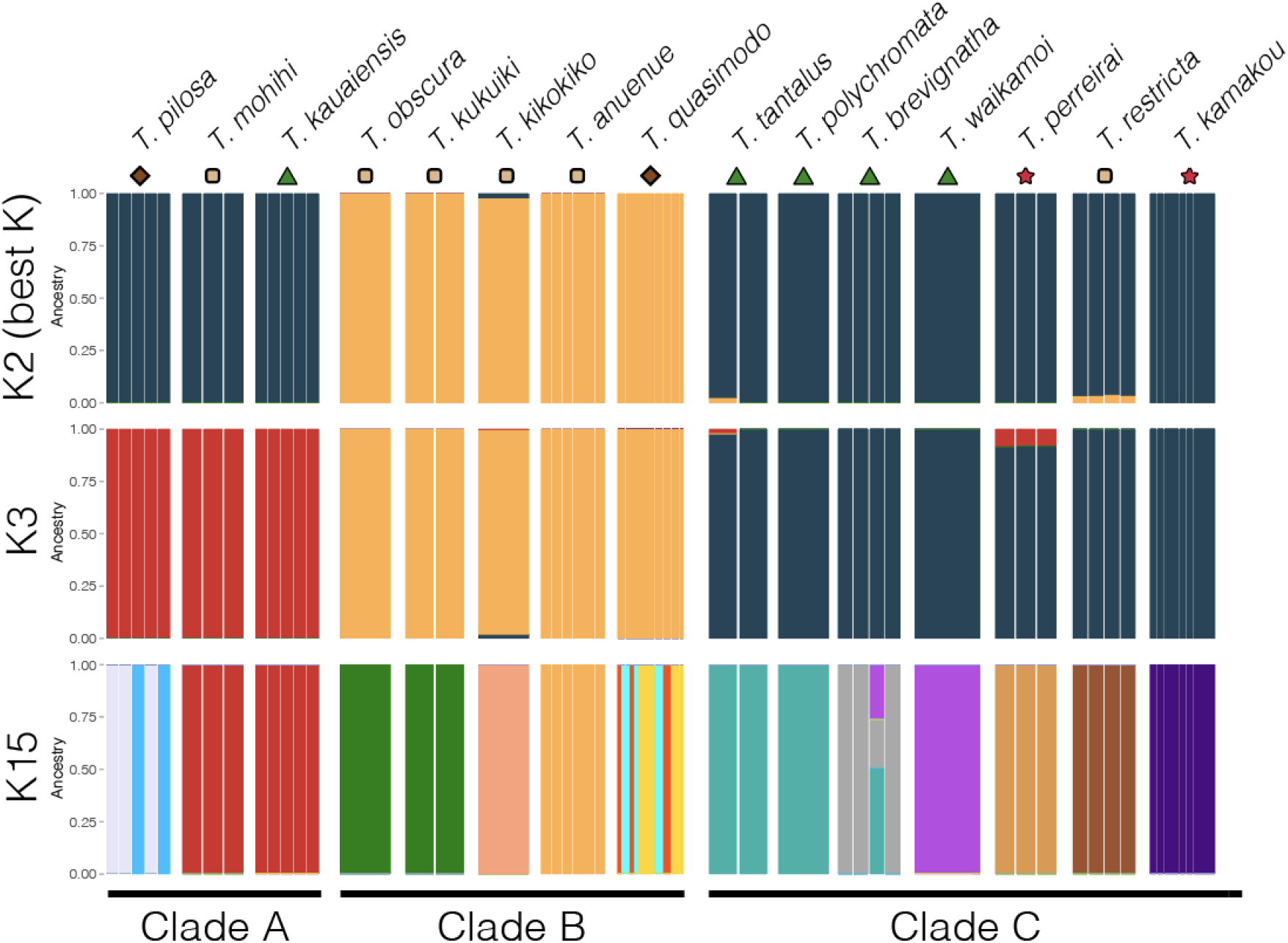
Structure of genetic diversity. NgsAdmix analysis with cluster (*K*) values of 2 (best *K*), 3 (major clades), and 15 (number of species).

### Excess allele sharing

Based on Patterson’s D statistics (Supplementary Figure 02) and F4-branch statistics, we found excess allele sharing between ecomorphs and within ecomorphs (Figure 4). We detected the largest excess in allele sharing between the Maroon ecomorph species (*T. perreirai* and *T. kamakou;* the trio P1 = *T. restricta;* P2 = *T. kamakou;* P3 = *T. perreirai* had a D-statistic of 0.15, Z-score 30.42 over >150,000 sites). In the Small Brown ecomorph, we also detected excess allele sharing, specifically between *T. restricta* and the remaining species (Figure 4; the trio P1 = *T. perreirai;* P2 = *T. restricta;* P3 = *T. obcura-kukuiki-kikokiko-anuenue* trio had a D-statistic of 0.13, Z-score of 8.21; >150,000 sites; Supplementary Table 03). No evidence for excess allele sharing was found between Green and Large Brown ecomorphs (Figure 4). When comparing ecomorphs, we find evidence of excess allele sharing between the lineages of clade A, namely between *T. pilosa* (Large Brown ecomorph), and *T. kauaiensis* and *T. mohihi* (Small Brown ecomorph); there is also excess of allele sharing between the Green ecomorph group (*T. tantalus, T. polychromata, T. brevignatha, T. waikamoi*) and one of the Maroon ecomorph species (*T. kamakou*). The Large Brown species *T. quasimodo* had excess allele sharing with the clade consisting of *T. restricta* (Small Brown ecomorph) and *T. kamakou* (Maroon ecomorph).

**Figure 4.**
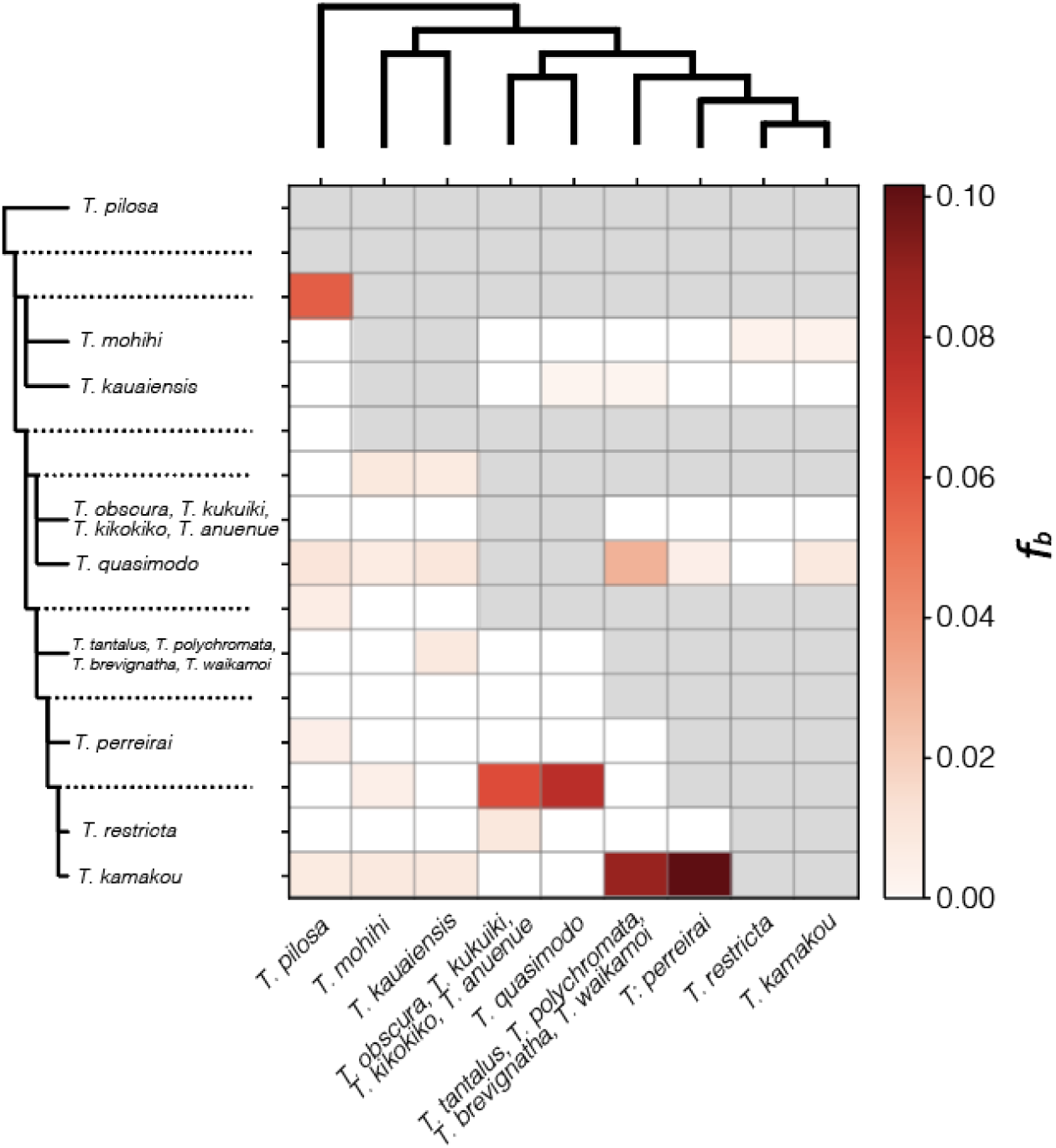
Excess allele sharing. F4-branch statistic plotted as a heatmap. The tree topology obtained by NgsDist and UCEs is plotted above and on the left every branch of the tree is displayed (including internal branches). Because the goal is to understand hybridization between ecomorphs, clades of closely related species belonging to the same ecomorph (i.e. monophyletic ecomorphs) are displayed.

The windowed-analyses of D-statistics and TWISST showed no particular pattern of introgression for *T. restricta, T. kamakou* and *T. perreirai*. Specifically, D-statistics of regions where melanization genes were found (see F_ST_ scans below) were not elevated. The TWISST analyses showed no specific tracks of hybridization in the largest three scaffolds (Supplementary Figures 3-5).

### F_ST_ scans

We ran F_ST_ comparisons for lineages with five or more individuals (Figure 5). The two within-ecomorph comparisons (i.e. Large Brown (a) *vs* Large Brown (b); Green (a) *vs* Green (b)) yielded no significant F_ST_ outliers, and this is likely attributed to the overall high F_ST_ between more distantly related species in the radiation (Large Brown (a) *vs* Large Brown (b) had an global F_ST_ of 0.60; Green (a) *vs* Green (b) had an global F_ST_ of 0.61). The remaining comparisons, displayed in Figure 5, all had overall F_ST_ between 0.31 (Green (b) *vs* Maroon) - 0.41 (Small Brown *vs* Maroon).

**Figure 5.**
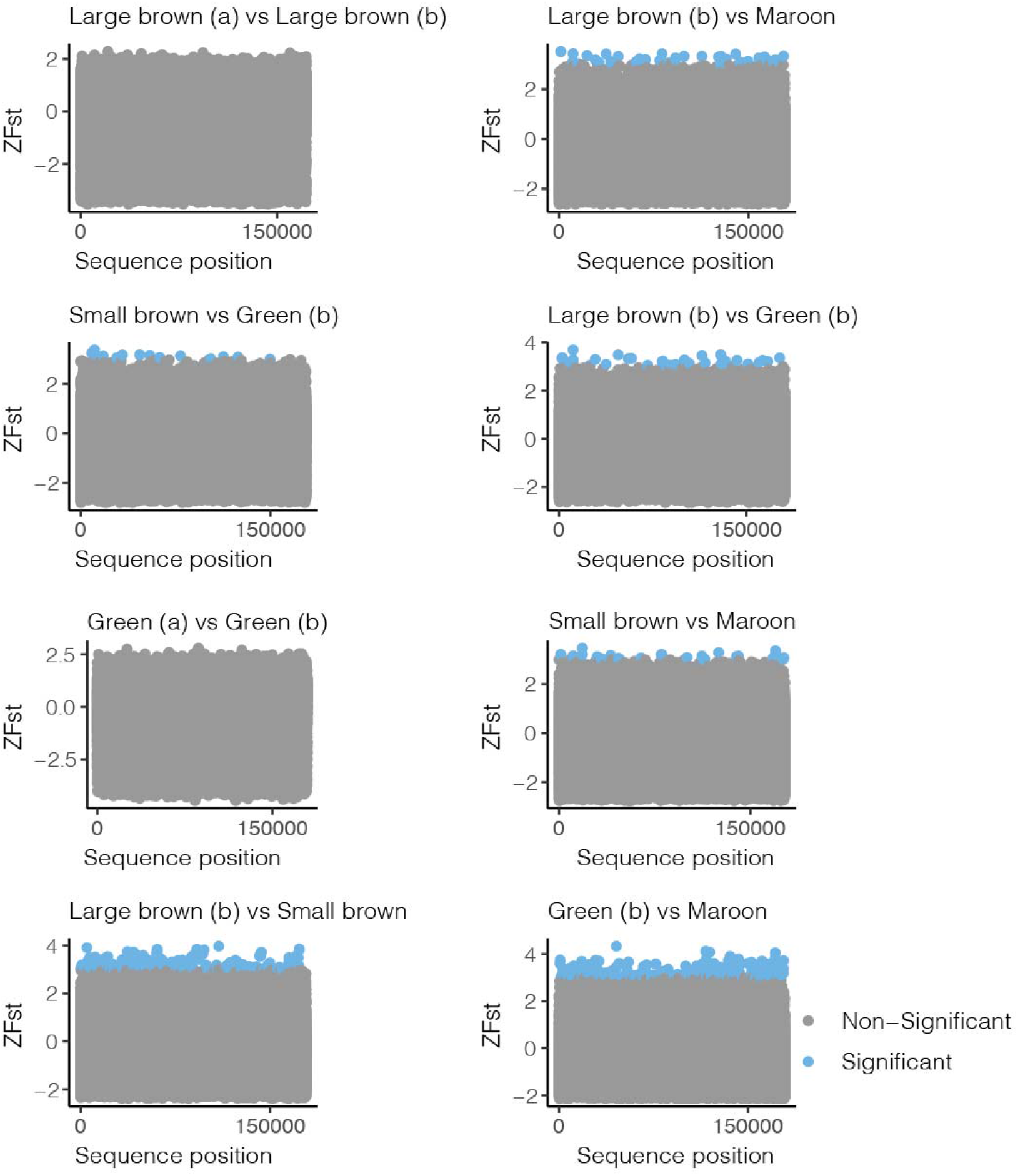
F_ST_ Scans of genomic divergence. For each pairwise comparison, we provide the sequence position and a Z-transformed F_ST_ value. Significant Z values are displayed in blue (Z > 3) and non-significant in grey. Large Brown (a) includes *T. pilosa* (5 individuals); Large Brown (b) includes *T. quasimodo* (16 individuals); Green (a) includes *T. kauaiensis* (5 individuals); Green (b) includes *T. tantalus, T. polychromata, T. brevignatha, T. waikamoi* (15 individuals); Small Brown includes *T. obscura, T. kukuiki, T. kikokiko, T. anuenue* (10 individuals); Maroon includes *T. kamakou* (9 individuals).

Several areas of divergence along the genome seem to be common in pairwise comparisons. Specifically, while significant F_ST_ can be driven by a highly divergent region in a single lineage, we found that multiple regions of the genome seem to be commonly differentiated. For example, we found twelve genomic regions where significant F_ST_ values were identified in different pairwise comparisons. As an example, the region between 10,000 - 15,000 bp on Scaffold-2,488 had F_ST_ outliers when comparing Large Brown (b) *vs* Green (b) and Small Brown *vs* Maroon.

The regions with significant genomic divergence yielded some interesting genes (Table 1; Supplementary Table 02). Notably, we found genes associated with circadian rhythms in all five comparisons, and genes associated with neuronal functions in four comparisons, and learning and memory formation in three comparisons (Table 1). Interestingly, we found one gene associated with diet (Starch digestion) and another with smell. Two genes were associated with longevity and life-span, and three genes were associated with growth or development (Table 1). Three of the comparisons included melanization-associated genes.

**Table 1.** Summary of genes in areas with high genomic divergence (F_ST_).

| Comparison | Gene | Function | References |
| --- | --- | --- | --- |
| Large Brown (b) vs Green (b) | NAT1 | Circadian rhythm | (Bradley et al. 2012) |
| Large Brown (b) vs Green (b) | ALT | Neuron feeding source | (Volkenhoff et al. 2015) |
| Large Brown (b) vs Green (b) | Dscam | Immune-response associated | (Nazario-Toole et al. 2018) |
| Large Brown (b) and Small Brown | Mal-B1 | Starch digestion | (Wu et al. 2016) |
| Large Brown (b) and Small Brown | vhaSFD | Longevity and life span | (Paaby and Schmidt 2009; Proshkina et al. 2015) |
| Large Brown (b) and Small Brown | Ank | Synaptic activity | (Koch et al. 2008) |
| Large Brown (b) and Small Brown | NAT1 and Gnpnat | Circadian rhythm | (Mattila and Hietakangas 2017) |
| Large Brown (b) and Small Brown | CG8483, and anon-W00118547.80 | Melanization-associated | (Bastide et al. 2016) |
| Large Brown (b) and Small Brown | CG8483 | Posterior development genes | (Ibrahim et al. 2013) |
| Large Brown (b) and Maroon | ND-B17 and ALT | Synapsis | (Ghosh 2013) |
| Large Brown (b) and Maroon | NAT1, MFS10, and Period circadian protein | Circadian rythm | (Curtin et al. 1995; Fernandez-Chiappe et al. 2021) |
| Large Brown (b) and Maroon | LIMK1 | Learning and memory formation | (E. A. Nikitina, A. V. Medvedeva, Yu. F. Dolgaya, L. I. Korochkin, G. V. Pavlova & E. V. Savvateeva-Popova 2012) |
| Green (b) and Maroon | CG18473 | Growth-regulation | (Hevia et al. 2017) |
| Green (b) and Maroon | Syp, CG8545, and Trpgamma | Synapses, neurojunctions, neurons and nervous-system related | (Guan et al. 2005; Liebl et al. 2006; Yang et al. 2017) |
| Green (b) and Maroon | LIMK1 | Learning and memory formation | (E. A. Nikitina, A. V. |
|  |  |  | Medvedeva, Yu. F.<br>Dolgaya, L. I.<br>Korochkin, G. V.<br>Pavlova & E. V.<br>Savvateeva-Popova<br>2012) |
| Green (b) and Maroon | MSF10,<br>Jhedup | Circadian behaviour<br>and rhythm | (Bloch et al. 2013;<br>Fernandez-Chiappe et<br>al. 2021) |
| Green (b) and Maroon | anon-WO0118547.8<br>0 | Melanization-associated | (Bastide et al. 2016) |
| Green (b) and Maroon | SNX3 | Development and<br>morphogenesis | (Zhang et al. 2011) |
| Green (b) and Maroon | RpS8 | Extention of lifespan | (Hoffmann et al.<br>2013) |
| Green (b) and Maroon | CG11471 | Smell-sensory<br>functions | (Anholt et al. 2001) |
| Small Brown and Maroon | anon-WO0118547.8<br>0 | Melanization-associated | (Bastide et al. 2016) |
| Small Brown and Maroon | Per | Circadian rhythm | (Curtin et al. 1995) |
| Small Brown and Maroon | LIMK1 | Learning and memory formation | (E. A. Nikitina, A. V. Medvedeva, Yu. F. Dolgaya, L. I. Korochkin, G. V. Pavlova & E. V. Savvateeva-Popova 2012) |

## Discussion

The three main findings of our work are: ***(i)*** The repeated evolution of *Tetragnatha* spiny-leg ecomorphs involves more than a single genomic-basis; ***(ii)*** The genomic-basis of ecomorph differentiation included genes associated with melanization, agreeing with previous ecological and morphological evidence which reported differences in colouration (Gillespie 2004; Kennedy et al. 2022); and ***(iii)*** pairwise scans of divergence uncovered evidence for a new axis of differentiation between ecomorphs involving neural and circadian rhythm changes.

### Evolutionary history of the *Tetragnatha* spiny-leg clade

Evidence of repeated evolution of ecomorphs comes from the paraphyletic and polyphyletic positioning of ecomorph species in the nuclear whole-genome phylogeny (Figure 2), and partitioning of genetic variation in admixture analyses (Figure 3). These analyses show that the spiny-leg radiation can be divided into three large clades. The first clade comprising the lineages in Kaua’i, the oldest island, and including three ecomorphs, namely Green (*T. kauaiensis*), Small Brown (*T. mohihi*), and Large Brown (*T. pilosa*). This clade is monophyletic and resolved with high confidence (bootstrap of 100) in the mitochondrial-genome tree (Figure 2), corroborating previous results from both nuclear and mitochondrial DNA (Pons and Gillespie 2003; Gillespie 2004). However, the whole-genome data suggests that *T. pilosa* is sister to the remaining radiation, grouping *T. kauaiensis* and *T. mohihi* as its own clade. The branches for *T. kauaiensis, T. mohihi* and *T. pilosa* are short, and this could indicate a lower substitution rate, higher missing data, incorrect attachment point of the outgroups, or root misplacement due to artefacts such as low-branch attraction. Lower substitution rate seems the most plausible of these scenarios as missing data can be discarded considering the filtering of the data (Supplementary Table 01). Attachment of outgroups follows the taxonomic knowledge of the group (Kennedy et al. 2022). Root misplacement due to NJ-algorithms, which can be prone to artefacts such as long-branch attraction (Susko et al. 2004), can be indirectly excluded as the UCE-based tree displays a similar topology for these branches and was done using ML. The nuclear-mitochondrial discordance pattern may result from hybridization (Figure 4) and from the fast diversification of the radiation, which can lead to an increase of rates of incomplete lineage sorting (Suh et al. 2015,Cerca et al. 2021b). Evidence for a fast diversification comes from the short branch and the low bootstrap support of the corresponding node (orange circle; Figure 2).

The second clade includes the widely distributed Large Brown species *T. quasimodo* and the Small Brown group including *T. obscura, T. kukuiki, T. kikokiko*, and *T. anuenue*, whose constituent species are present on the islands of Hawai⍰i, O⍰ahu, Maui and Moloka⍰i. The nuclear and the mitochondrial lineages show topological congruence (Figure 2B). The third clade includes a group of Green ecomorph species (*T. tantalus, T. polychromata, T. waikamoi*, and *T. brevignatha*) which are distributed on several islands (Hawai⍰i, O⍰ahu, Maui), two separate lineages with Maroon ecomorphs (*T. kamakou* from Moloka⍰i and Maui; *T. perreirai* from O⍰ahu), and a Small Brown ecomorph species (*T. restricta*). In this clade, we found evidence for nuclear-mitochondrial disagreements (Figure 2 A-B) and likely hybridization (Figure 4). Most prominently, part of the group comprised by Green ecomorphs, namely *T. tantalus* and *T. polychromata*, and the Maroon *T. perreirai*, nests as sister to the second clade in the mitochondrial tree, rendering this clade paraphyletic in the mitochondrial dataset.

The Green ecomorph species *Tetragnatha macracantha* (Haleakalā volcano in Maui, and Lana’i) was not included due to the lack of specimens. However, based on previous studies (Gillespie 2004; Cotoras et al. 2018) it clusters with the other Green ecomorph species present in Maui (*T. waikamoi* and the Maui population of *T. brevignatha*). Therefore, the described patterns most likely also include this species. Similarly, *Tetragnatha kukuhaa*, a Small Brown from the Big island is not-included, but it has been previously suggested as a sister to *T. obscura* (Gillespie 2004; Kennedy et al. 2022).

### Genomic basis of ecomorph evolution

We find evidence that the evolution of ecomorphs has happened repeatedly, in agreement with previous works (Gillespie 2004), and extend these findings by finding evidence that multiple genomic sources underlie repeated phenotypic evolution. Specifically, we find evidence for excess allele sharing, which usually results from hybridization. The strongest signal for hybridization in the dataset, where as much as 10% of the whole-genome variants may be introgressed, occurs from *T. perreirai* to *T. kamakou*, the two species belonging to the Maroon ecomorph (Figure 4). This evidence is further corroborated by the nuclear-mitochondrial disagreements (Figure 2). Noticeably, *T. perreirai* and *T. kamakou* are not sister species in any of the phylogenetic analysis carried by us (Figure 1; Supplementary Figure 01) and do not currently overlap geographically (*T. perreirai* occurs on the island of O⍰ahu, *T. kamakou* on Maui and Moloka⍰i). Several scenarios that could have led to the lack of monophyly of this group in the face of hybridization.

First, introgression may have occurred between *T. perreirai* and an ancestral lineage that was present on Maui or Moloka⍰i. This introgression event may have carried adaptive alleles associated with the Maroon ecomorph, and opened the Maroon-niche to the admixed lineage, which is now *T. kamakou*. This event of introgression may have been subtle as our exploration of genomic windows for the largest scaffolds (TWISST) and for genomic windows (Fd ratio) where we found genes associated with melanization (CG8483, anon-WO0118547.80), yielded no evidence for introgression. It is possible that we missed signals of introgression due to the sparsity of low-coverage data along the genome, since missing data may blur genomic patterns and signals (Cerca et al. 2021b). Additionally, we cannot exclude recurrent ancestral hybridization, and this may blury the distinction between hybridization and standing genetic polymorphism (Ferreira et al. 2021). In any case, the admixture analyses suggested no shared ancestry tracks between these three species, even at *K* = 15, and this indicates that signatures of introgression may have been diluted through time.

The second scenario is that the Maroon phenotype evolved first in *T. kamakou*, and was introgressed to O⍰ahu, leading to the evolution of *T. perreirai*. This alternative scenario implies a back colonization of O’ahu, which is older than Maui Nui. At this stage, we cannot confidently ascertain any of these scenarios, and further data including more genomes, higher coverage, and analyses such as demographic simulations are necessary. We speculate that the combination of ancestral bouts of introgression, fast diversification creating incomplete lineage sorting, changes in population size, population-extinctions as a result of island-cycles, as observed in other adaptive radiations (Meier et al. 2018; De-Kayne et al. 2022), limits our capacity to understand the evolution of this group.

We also find evidence of excess allele sharing between members of the Small Brown ecomorph. Namely, there is a signal of hybridization between *T. restricta, T. mohihi*, and the clade comprising 4 Small Brown ecomorph species (*T. anuenue, T. kukuiki, T. obscura, T. kikokiko*). This signal is particularly clear from the Patterson’s D analysis (Supplementary Figure 02), but less clear from the F4 ratio test (Figure 4). This mismatch is likely explained by the different topologies underlying both tests. Two scenarios can explain these results. First, considering that we find evidence for excess lineage sorting between lineages on different islands, it can be speculated that dispersal may have occurred at some time in the past, leading to hybridization. Second, hybridization is not the only cause leading to excess allele sharing and this can confound Patterson’s D and F4 -ration analyses. For example, the occurrence of speciation between lineages with different population sizes may cause asymmetries that lead to elevated Patterson’s D statistics (smaller populations will have less variants due to the effect of drift). Similarly, if multiple alleles underlie the evolution of an ecomorph, there may be imbalances in allelic variation along the genome, confounding patterns of allele excess. While scenarios of population-size differences and multi-alleles are hard to rule out with low-coverage data, they are unlikely as we also observe mitochondrial-nuclear discordances. Our results contribute to the growing body of work showing that introgression can be a potent force in the passage of adaptive alleles from one lineage to another (Meier et al. 2017; Marques et al. 2019; Sowersby et al. 2021) and can thereby drive repeated phenotypic evolution in the context of an adaptive radiation.

Despite the evidence of hybridization in the spiny-leg lineage, hybridization alone is not sufficient to explain the repeated evolution of every ecomorph. Specifically, we do not find evidence for excess allele sharing between the two Green ecomorph clades (*T. kauaiensis* and the clade comprising the lineages *T. brevignatha, T. waikamoi, T. tantalus, T. polychromata*) and a weak signal of hybridization between *T. pilosa and T. quasimodo*. The lack of hybridization within the Green ecomorph is consistent with previous studies, which found no hybridization between Green ecomorph species (Cotoras et al. 2018). The repeated evolution of some ecomorphs may have occurred by either *de novo* mutation or ancestral genetic variation (Barrett and Schluter 2008). Nonetheless, distinguishing between these scenarios benefits from the study of sweeps and coalescence of variants (Barrett and Schluter 2008; Lee and Coop 2017), which would require higher coverage and wider population sampling obtained by us. Despite these limitations, several pieces of evidence may point towards a role of repeated phenotypic evolution of Green and the Large Brown ecomorphs through standing genetic variation. First, the repeated recruitment of variation may be more likely than repeated *de novo* mutations producing the same phenotype (Barrett and Schluter 2008). Second, as discussed above, the phylogenetic reconstruction includes some short branches, which drives higher rates of incomplete lineage sorting, translating into standing genetic variation passing down the lineages. Third, we found 12 genomic regions that were repeatedly significantly-diverged in pairwise F_ST_ comparisons. However, we are careful not to exclude the existence of *de novo* mutation in the radiation and in specific genomic regions, which could be hotspots of mutation with some adaptive value; e.g. (Xie et al. 2019).

Because adaptive introgression seems to play a role in the evolution of some ectomorphs (Small Brown and Maroon), but not in others (Green and Large Brown), this opens the question as to how easy it becomes to re-evolve certain traits without hybridization. In other words, is the transfer of some key genes a strict requirement for the appearance of the Small Brown and Maroon ecomorphs? If that were to be the case, this could explain why the Maroon ecomorph is entirely absent from Lana’i and the Big Island (Gillespie 2004).

### Ecological and genomic drivers of ecomorph evolution

Spiny-leg *Tetragnatha* spiders are largely confined to mid-elevation wet and mesic forests on Hawaiian volcanoes (1,200-1,800 meters) and diversification has occurred largely within this environment (Hiller et al. 2019). This suggests that, in addition to the physical barrier between islands imposed by the ocean, the lowland area may act as a strong isolating barrier, as has been shown for many taxa in more stable environments (Janzen 1967). The spiders’ forest habitats have mosaic distributions, and it is likely that the separation between them has triggered a dynamic interplay between natural selection in response to micro-habitat availability and allopatric divergence (Vandergast et al. 2004; Roderick et al. 2012; Cotoras et al. 2018). In particular, ecomorph divergence is likely due to allopatric establishment on different islands and volcanoes, followed by secondary contact, competition between ecologically similar species and hence the accumulation of divergence (Schluter 2000; Cotoras et al. 2018). Dispersal and secondary contact of diverged populations may have led to a macroevolutive ecological character displacement and the evolution of particular ecomorphologies. This scenario is consistent with the evidence for hybridization at various points of the radiation, which suggests that dispersal may have occurred multiple times across the phylogeny, leading to the introgression of variants.

A major goal of adaptive radiation research is to disentangle the ecological drivers of species formation (speciation) and adaptation to the environment (local adaptation). Previous research has suggested that ecomorph-colouration has been associated with selective pressure from predators (Gillespie 2004). In the current study, F_ST_ comparisons uncovered melanization genes in regions of genomic divergence, which agree with the natural history observations of the group and the colour-characterization of the ecomorphs. Interestingly, we also find evidence for genes associated with circadian rhythms, neuronal and synaptic activity genes, life span, learning and memory formation, which can be seen as an unexplored axis of physiological and ecological differentiation. Specifically, these genes may indicate that ecomorph-evolution may involve more phenotypic changes than shifts in coloration and micro-habitat use. For example, circadian rhythm genes are found across the tree of life and are important for a wide array of functions, from allowing organisms to synchronise with their immediate environment to regulating reproductive activity, and it has been suggested that they play an important role in environmental adaptation (Yerushalmi and Green 2009). Finally, the evidence for genomic differentiation in developmental genes may suggest that shifts in developmental stages may have an important function in the *Tetragnatha* spiny-leg adaptive radiation. This is consistent with previous work which shows that the initial appearance of the Maroon ecomorph is associated with developmental shifts in coloration on the oldest island of Kauai (Brewer et al. 2014), by identifying several genes under selection likely responsible for changes in coloration and developmental processes – reticulon nogo and apolipoprotein d.

## Conclusion

The whole-genome sequencing of *Tetragnatha* spiny-leg ecomorphs showed that the genomic basis of repeated ecomorph evolution is multifarious even in closely related species, as some ecomorphs likely arose through hybridization (Small Brown, Maroon), while others likely arose by shared standing genetic variation or *de novo* mutation (Green, large Brown). We also found that ecomorph evolution in the spiny-legs may go beyond their colouration, as regions with high genomic divergence consistently pointed to genes associated with other potential ecological axes of differentiation, such as learning and memory formation, circadian rhythms, and developmental shifts. However, it is likely that a complex evolutionary history involving population extinctions, population structuring and ghost-lineages blurry the reconstruction and interpretation of evolutionary patterns.

## Author Contributions

JC, THS, RGG obtained funding. JC and RGG designed the study. RGG, DDC, SK, HK, AJW provided samples. JC, LH generated the data. JC analysed the data with contributions on scripting and interpreting the data from CGS, VCB, JM, DDC, JML, CFP and DD. JC drafted a first manuscript, and all the authors contributed to the writing and clarity of the manuscript.

## Data Accessibility

Data is being made available through ENA XXX. Code is at github José.

## Acknowledgements

The authors would like to acknowledge a large number of people and institutions that collaborated at different stages of this research. The fieldwork in Maui Nui was supported by Timothy Bailey and William Roderick. JC is grateful to Mike Martin for access to a super computer cluster, Mark Ravinet for his constant mentorship and LD scripts, and Michael Matschiner for comments on the manuscript. We are grateful to Jun Ying Lim for providing samples.

The permit processing and access to different reserves and private land was possible thanks to Pat Bily (TNC Maui), Lance DaSilva (DOFAW Maui), Danae Dean (Kahoma Ranch), Charmian Dang (NAR), the late Betsy Gagne (NAR), Elizabeth Gordon (HALE), Paula Hartzell (Lana⍰i Resorts, LLC), Pomaika⍰i Kaniaupio-Crozier (Maui Land and Pineapple), Cynthia King (DLNR), Peter Landon (NAR Maui), Russell Kallstrom and Ed Misaki (TNC Moloka⍰i) and Joe Ward (Maui Land and Pineapple).

JC, RG, THS were supported by a Peder Sather grant, DDC was supported by a Fulbright/CONICYT doctoral fellowship, Integrative Biology department and the Graduate Division of UC Berkeley, the Margaret C. Walker fund (Essig Museum of Entomology) and a Humboldt postdoctoral fellowship. Fieldwork was funded by NSF grants DEB 1241253 and DEB 1927510 to RG.

## Notes

### Competing Interest Statement

The authors have declared no competing interest.

